# Cisplatin exposure dysregulates pancreatic islet function in male mice

**DOI:** 10.1101/2024.09.25.614996

**Authors:** Lahari Basu, Kelsea S. McKay, Myriam P. Hoyeck, Lili Grieco-St-Pierre, Kayleigh R.C. Rick, Emilia Poleo-Giordani, Evgenia Fadzeyeva, Erin E. Mulvihill, Jan A. Mennigen, Jennifer E. Bruin

## Abstract

Cancer survivors have an increased risk of developing new-onset Type 2 diabetes compared to the general population. Moreover, patients treated with cisplatin, a commonly used chemotherapeutic agent, are more likely to develop metabolic syndrome and Type 2 diabetes compared to age- and sex-matched controls. Insulin-secreting beta cells—located within pancreatic islets—are critical for maintaining glucose homeostasis, and dysregulated insulin secretion is central to Type 2 diabetes pathophysiology. Surprisingly, the impact of cisplatin treatment on pancreatic islets has not been reported. In this study, we aimed to determine if murine islet function is adversely affected by direct or systemic exposure to cisplatin. *In vitro* cisplatin exposure to male mouse islets profoundly dysregulated insulin release, reduced oxygen consumption, and altered the expression of genes related to insulin production, oxidative stress, and the Bcl-2 family. *In vivo* cisplatin exposure led to sustained hypoinsulinemia and hypoglycemia in male mice. Pancreas tissues from cisplatin-exposed male mice showed increased proinsulin accumulation and expression of DNA-damage markers in beta cells, but no change in average islet size or % insulin^+^ area per islet. Our data suggest both direct and systemic cisplatin exposure cause acute defects in insulin secretion and may have lasting effects on islet health in mice.

## Introduction

Diabetes incidence is rising rapidly, marking this condition as a major global health concern(1). Glucose homeostasis is managed by endocrine cells within the pancreas, known as the islets of Langerhans. Type 2 diabetes (T2D), which accounts for approximately 90% of all diabetes diagnoses, is characterized by chronic hyperglycemia, impaired insulin secretion from pancreatic beta cells, and defects in peripheral insulin action(1,2).

Numerous studies have shown that cancer survivors—irrespective of age, treatment, and type of cancer—have an increased risk of developing new-onset T2D compared to the general population(3–5). Moreover, cancer survivors often develop T2D earlier than control populations(6,7). Sylow *et al.*(8) recently reported that out of 28,000 cancer survivors, those with new-onset T2D had a 21% higher all-cause mortality compared to cancer survivors without T2D. Therefore, cancer survivors are not only more vulnerable to diabetes than the general population, but those who develop diabetes have higher rates of mortality than cancer patients without secondary diabetes.

Cisplatin is a platinum-based chemotherapeutic agent used to treat a wide variety of cancers, including lung, ovarian, and colorectal cancers(9). Cisplatin can enter cells through passive diffusion or through the copper transporter 1 protein(10), a protein that has not been well characterized in islets. Once cisplatin enters the cell, either one or both chlorine groups are replaced by water molecules, effectively becoming aquated and biologically active(11). Aquated cisplatin is highly electrophilic, allowing it to bind to neutrophilic centers on purine residues in nuclear and mitochondrial DNA, causing the crosslinking of DNA, which induces DNA damage responses and apoptotic pathways(11). Cisplatin also increases the production of reactive oxygen species (ROS) and leads to oxidative stress and mitochondrial deterioration(11–13). This, in turn, leads to increased cell senescence and cell death(11).

Unfortunately, the cytotoxic effects of cisplatin are not limited to only cancerous cells. Cisplatin treatment has been linked to acute toxicities and severe off-target effects, including nephrotoxicity and neurotoxicity(14). There is also evidence linking cisplatin treatment and metabolic complications. Testicular cancer patients treated with cisplatin had increased odds of developing metabolic syndrome compared to patients treated with surgery or radiotherapy and the general population(15). In a group of 219 patients who received cisplatin-based chemotherapy for head and neck cancer, 5% developed diabetes during their treatment period(16). This is almost twice the global rate of diabetes prevalence at the time of the study(17).

The potential off-target effects of cisplatin on pancreatic islets are currently unknown. Given the known roles of DNA damage, mitochondrial dysfunction, and oxidative stress in beta cell dysfunction during T2D pathogenesis(18–20), we hypothesized that beta cells would be particularly susceptible to the off-target effects of cisplatin. Moreover, the limited regenerative capacity of beta cells(21) would likely prevent recovery of the beta cell population following injury and contribute to increasing risk of T2D in cancer survivors who receive cisplatin treatment. In our study, we assessed the effects of cisplatin on beta cell function by treating isolated male mouse islets with cisplatin *in vitro* and exposing male mice to cisplatin *in vivo*. We found that *in vitro* cisplatin exposure profoundly dysregulated insulin secretion and mitochondrial function in isolated mouse islets. Cisplatin-exposed mice had reduced plasma insulin levels and increased markers of defective proinsulin processing and DNA damage in beta cells at 6 weeks post-exposure compared to vehicle-exposed mice. Overall, our data suggest that cisplatin exposure impairs beta cell function in mice.

## Methods

### Mouse islet isolation and culture

Male C57Bl/6N mice were euthanized by cervical dislocation. Pancreata were inflated via common bile duct injection with collagenase (1000 IU/mL; Sigma-Aldrich, #C7657) dissolved in Hanks’ balanced salt solution (HBSS; 137 mM NaCl, 5.4 mM KCl, 4.2 mM NaH_2_PO_4_, 4.1 mM KH_2_PO_4_, 10 mM HEPES, 1 mM MgCl_2_, 5 mM anhydrous dextrose, pH 7.2) and excised. Pancreas tissues were incubated at 37°C for approximately 10 minutes 30 seconds then vigorously agitated, after which the collagenase reaction was immediately ceased by adding cold HBSS with 1 mM CaCl_2_. The digested pancreas tissues were then washed 3 times with HBSS with CaCl_2_, and islets were separated and purified using a Histopaque gradient (Sigma-Aldrich, #10771). Islets were then filtered through a 70 μm cell strainer, resuspended in RPMI 1640 1X (Gibco, #11875093 or Wisent Bioproducts, #350-000-CL) supplemented with 10% (vol./vol.) fetal bovine serum (FBS; Sigma-Aldrich, #F1051-500ML) and 1% (vol./vol.) penicillin-streptomycin (Corning, #30-002-Cl or Gibco, #15140-122-100), and handpicked under a dissecting scope to >95% purity. Islets were incubated in complete RPMI overnight at 37°C and 5% CO_2_ before *in vitro* chemical exposure.

### *In vitro* cisplatin exposure protocol

Following an overnight incubation in complete RPMI mouse islets were handpicked, and half of the islets from each biological replicate were incubated with complete RPMI containing either 10 μM cisplatin (Sigma-Aldrich, #232120-50MG) and the other half were incubated with saline (vehicle control) for up to 48 hours. The dose of cisplatin was chosen after a review of the literature(13,22) and a LIVE/DEAD^TM^ assay was conducted to ensure this dose did not cause significant cell death compared to vehicle-exposed cells over the course treatment. Media was refreshed after 24 hours with complete RPMI containing vehicle or cisplatin.

### LIVE/DEAD^TM^ assay

To assess cell viability, 50 islets per replicate (n = 2 technical replicates per mouse; n = 6 mice/group) were handpicked after 6, 24, and 48 hours of chemical exposure, and washed with pre-warmed (37°C) PBS (Sigma, #D8662). Islets were incubated in 400 μL Accutase (Corning, #25-058-CL) at 37°C for approximately 4 minutes, with tituration every 2 minutes to disperse islets into single cells. Accutase was neutralized with an equal volume of complete RPMI media to stop digestion. Cells were washed once with PBS then incubated with complete RPMI media containing 0.5 μM Hoescht (Thermo Fisher, #62249), 1.25 μM Calcein (Invitrogen, #L3224), and 0.5 μg/mL propidium iodide (PI; Invitrogen, #P21493) dyes. From each single-cell suspension, 2 technical replicates of 200 μL of cells were transferred to a black 96-well optical-bottom plate and incubated at room temperature for 30 minutes with 5% CO_2_. An Axio Observer 7 microscope was used to image 10% of each well immediately after incubation. The number of calcein^+^ cells (live) and PI^+^ cells (dead) were quantified using Zen Blue 2.6 software (Carl Zeiss). The percentage of PI^+^ cells was calculated as [(number of PI^+^ cells imaged in well) / (total number of cells imaged in well) × 100].

### Glucose-stimulated insulin secretion assays

To assess static glucose-stimulated insulin secretion (GSIS), 25 islets per replicate (n = 1-3 technical replicates/mouse; n = 7-8 mice/group) were handpicked after a 48-hour chemical exposure period and washed with pre-warmed (37°C) Krebs-Ringer bicarbonate HEPES buffer (KRBH) without glucose (115 mM NaCl, 5 mM KCl, 24 mM NaHCO_3_, 2.5 mM CaCl_2_, 1 mM MgCl_2_, 10 mM HEPES, 0.1% (wt/vol) BSA, pH 7.4). Islets were then incubated in KRBH with 2.8 mM glucose (low glucose; LG) for a 1-hour pre-incubation at 37°C with 5% CO_2_ after which the supernatant was discarded. Following this, islets underwent sequential 1-hour incubations in LG KRBH and KRBH with 16.7 mM glucose (high glucose; HG) at 37°C with 5% CO_2_. After each incubation, samples were centrifuged and supernatant was collected and stored at −30°C. To measure insulin content, islets were immersed in an acid-ethanol solution (1.5% (vol./vol.) HCl in 70% (vol./vol.) ethanol) at 4°C overnight, then neutralised with equal volume 1 M Tris base and stored at −30°C until analysis by ELISA.

To assess GSIS dynamically, 70 islets per mouse (n = 4 mice/group) were handpicked after 48 hours of chemical exposure and washed with pre-warmed PBS. Islets were then loaded in Perspex microcolumns between two layers of acrylamide-based microbeads (Biorep Technologies, #PERI-BEADS-20). Islets were perfused for 48 minutes at a rate of 100 μL/min with LG KRBH to equilibrate the islets. The islets were then perfused with LG KRBH for 8 minutes, HG KRBH for 32 minutes, LG KRBH for 20 minutes, KRBH with 30 mM KCl for 20 minutes, and LG KRBH for 16 minutes. Samples were collected every 2 minutes (200 μL total/sample). Islets and perifusion solutions were kept at 37°C throughout the perifusion run using the built-in temperature-controlled chamber while the collection plate was kept at 4°C using the built-in tray cooling system. Samples were stored at −80°C until analysis. Insulin concentrations in both static GSIS and perifusion samples were measured by rodent insulin chemiluminescence ELISA (ALPCO, #80-INSMR-CH10).

### Oxygen consumption analysis

Islet respiration was quantified using the Seahorse XFe24 Analyzer (Agilent Technologies). To assess mitochondrial function, 70 islets per mouse (n = 4 mice/group) were handpicked after 48 hours of chemical exposure and washed with Seahorse XF RPMI media (Agilent Technologies, 103576, pH 7.4) supplemented with 2 mM sodium pyruvate, 2 mM L-glutamine, 2.8 mM D-glucose, and 1% (vol./vol.) FBS. Islets were then plated in a 24-well islet capture plate (Agilent Technologies, #103518-100) coated with poly-d-lysine (Sigma-Aldrich, #P7280) and incubated at 37°C and 0% CO_2_ for 1.5 hours. Media was refreshed following the incubation before loading the plate into the XFe24 Analyzer. Basal oxygen consumption rate was measured for approximately 35 minutes. After this, wells were exposed to sequential injections of 16.7 mM glucose for 6 cycles, 2.5 μM oligomycin for 8 cycles, 3 μM carbonyl cyanide-p-trifluoromethoxyphenylhydrazone (FCCP) for 5 cycles, and finally a combination of 3 μM rotenone and 3 μM antimycin A for 6 cycles. With each cycle, the solutions were mixed for 3 minutes, samples were allowed to rest for 2 minutes, and oxygen consumption was measured for 3 minutes. The following parameters of mitochondrial function were calculated as per Table 1.

**Table 1.**
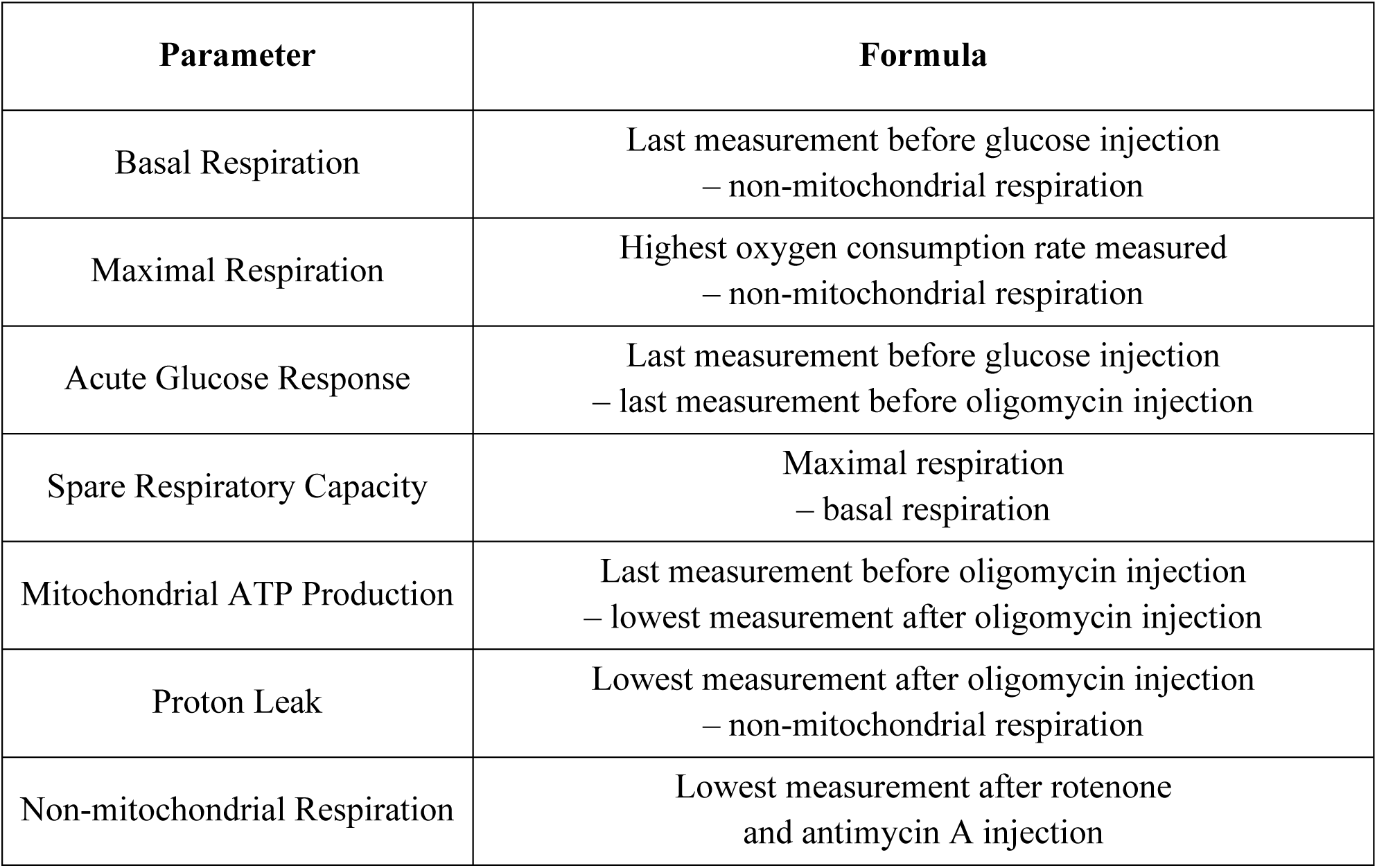
Formulas of mitochondrial function parameters.

| Parameter | Formula |
| --- | --- |
| Basal Respiration | Last measurement before glucose injection<br>– non-mitochondrial respiration |
| Maximal Respiration | Highest oxygen consumption rate measured<br>– non-mitochondrial respiration |
| Acute Glucose Response | Last measurement before glucose injection<br>– last measurement before oligomycin injection |
| Spare Respiratory Capacity | Maximal respiration<br>– basal respiration |
| Mitochondrial ATP Production | Last measurement before oligomycin injection<br>– lowest measurement after oligomycin injection |
| Proton Leak | Lowest measurement after oligomycin injection<br>– non-mitochondrial respiration |
| Non-mitochondrial Respiration | Lowest measurement after rotenone<br>and antimycin A injection |

### Quantitative real time PCR

A subset of islets was handpicked at 6, 24, and 48 hours after chemical exposure and stored in buffer RLT with 1% 2-mercaptoethanol. RNA was extracted using the RNeasy Micro Kit (Qiagen, #74004) as per manufacturer’s instructions. DNase treatment was performed prior to cDNA synthesis with iScript gDNA Clear cDNA synthesis Kit (Bio-Rad, #1725035). Following this, qPCR was performed using SsoAdvanced Universal SYBR Green Supermix (Bio-Rad, #1725271) and run on a CFX384 (Bio-Rad). All targets were run alongside “no reverse transcriptase” and “no cDNA template” controls. *Ppia* was used as the reference gene given its stable expression under both control and treatment conditions. Data were analyzed using the 2^-ΔΔCt^ method. Primer sequences are listed in Table 2.

**Table 2.**
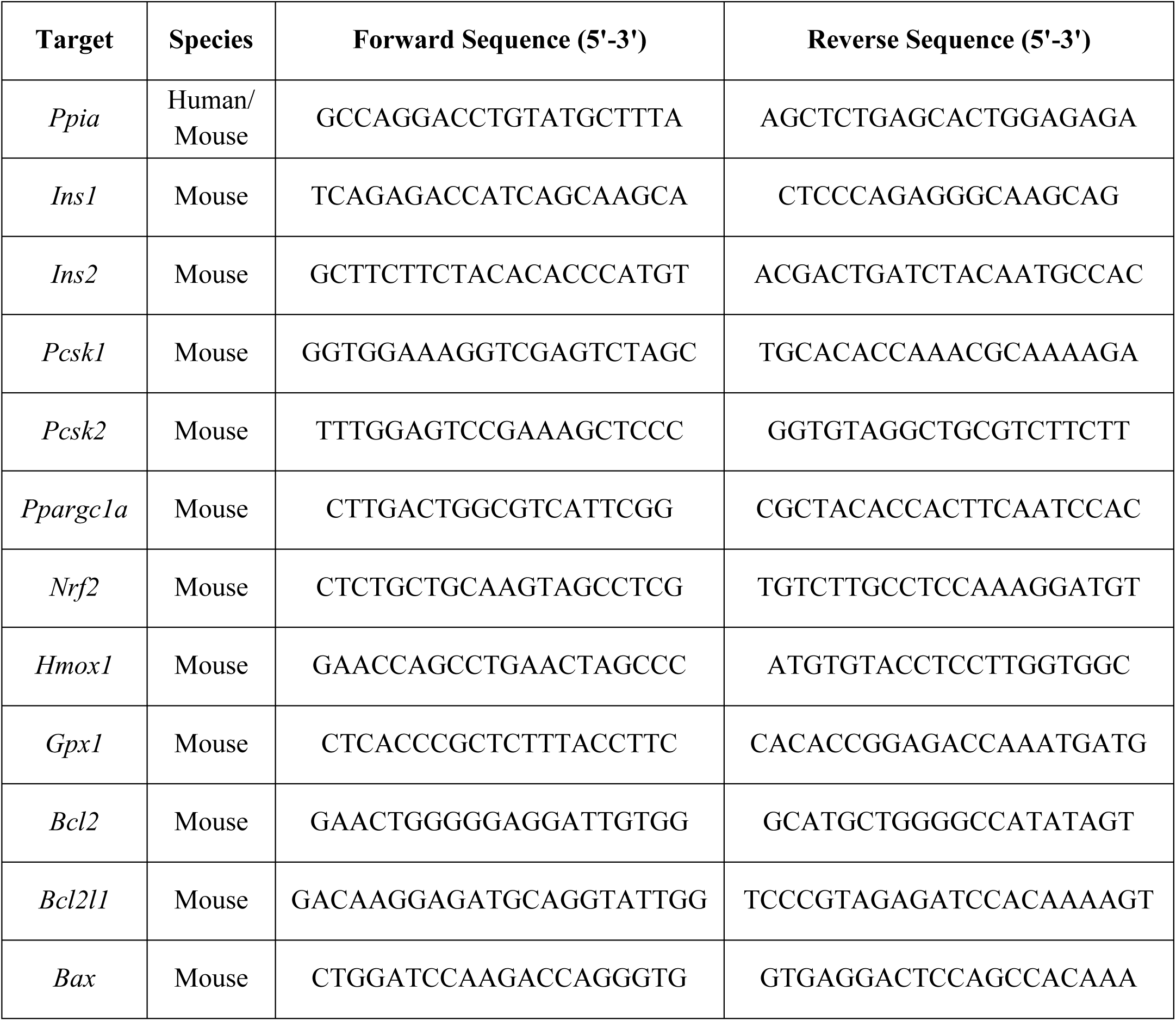
Primer sequences for qPCR.

### *In vivo* cisplatin exposure protocol

Male DBA/2 mice, between 6-8 weeks old (Charles River Laboratories) were maintained on a 12-hour light/dark cycle with *ad libitum* access to a standard rodent chow diet (Harlan Laboratories, Teklad Diet #2018) and water. All experiments were approved by Carleton University and University of Ottawa Animal Care Committees and carried out in accordance with Canadian Council on Animal Care guidelines. Prior to starting experimental protocols, animals were randomly assigned to treatment groups, and groups were matched for body weight and blood glucose, to ensure that these variables were consistent between groups.

As outlined in Figure 5A, mice received intraperitoneal (ip) injections of 0.2 mL 0.9% saline (vehicle control; n = 10) or 2 mg/kg cisplatin (Pfizer Canada, #02126613; n = 10) every other day over the course of 14 days, for a total of 7 injections. Although this dose is low relative to dosages used in clinical practice(23–25), this dosing protocol was chosen based on similar experimental protocols reported in the literature that showed induction of cisplatin-induced toxicity with less than 20% body weight loss(26–28).

### *In vivo* metabolic assessments

All metabolic analyses were performed in conscious, restrained mice, and blood samples were collected from the saphenous vein using heparinized microhematocrit tubes (Fisherbrand, #22-362-566). Blood glucose levels were measured using a OneTouch Verio Flex® glucometer (LifeScan). Body weight and blood glucose were measured following a 4-hour morning fast once a week throughout the study. For all metabolic tests, time 0 indicates the blood sample was collected prior to the administration of glucose or insulin.

For glucose tolerance tests (GTTs) conducted at 1- and 5-weeks post-exposure, mice received an ip injection of 2 or 3 g/kg glucose, respectively, following a 6-hour morning fast. Blood glucose levels were measured immediately prior to glucose-injection (referred to as 0 minutes post-glucose injection) and at 15, 30, 60, and 90 minutes post-glucose injection; blood samples were collected in heparinized tubes at 0, 15, and 30 minutes following glucose-injection and spun at 7000 rpm for 9 minutes to collect serum. Serum was used to measure plasma insulin levels by ultrasensitive rodent insulin ELISA (ALPCO, #80-INSMSU-E10). During an insulin tolerance test (ITT) conducted 4 weeks post-exposure, mice received an i.p. bolus of 1.15 IU/kg insulin (Novo Nordisk, Novolin ge Toronto, #02024233) following a 4-hour morning fast. Blood glucose was measured immediately before the insulin injection (0 minutes), then at 15, 30, 60, and 90 minutes post-insulin injection. For each metabolic assessment, mice from different treatment groups were randomly distributed throughout the assessment to ensure that the timing of blood collection was not a factor in our analysis. All mice received 0.2 mL saline at the end of each metabolic assessment for fluid replenishment.

### Immunofluorescence staining and image quantification

At 6 weeks post-exposure, pancreas tissues were harvested from mice and stored in 4% paraformaldehyde (PFA) for 24 hours, followed by long-term storage in 70% (vol./vol.) ethanol (n = 5 mice/group). PFA-fixed pancreas tissues were processed and paraffin-embedded by the University of Ottawa Heart Institute Histology Core Facility. Immunofluorescent staining was performed as previously described(29).

The following primary antibodies used in this study: rabbit anti-insulin (Cell Signaling, #C27C9, 1:200), mouse anti-glucagon (Sigma-Aldrich #G2654, 1:250), mouse anti-proinsulin (Developmental Studies Hybridoma Bank, #GS-9A8-s, 1:50), mouse anti-insulin (Cell Signalling, #L6B10, 1:250), rat anti-p21 (Abcam, #Ab107099, 1:1000), and rabbit anti-gamma H2AX (Abcam, #Ab1174, 1:7000). The following secondary antibodies were used: goat anti-rabbit IgG (H+L), Alexa Fluor 594 (Invitrogen, #A11037, 1:1000), IgG (H+L), Alexa Fluor 488 (Invitrogen, #A11029, 1:1000), and goat anti-rat IgG (H+L), Alexa Fluor 594 (Invitrogen, #A11007, 1:1000).

For islet morphology quantification, a minimum of 5 islets per mouse were imaged with an Axio Observer 7 microscope and the average of all islet measurements was reported for each biological replicate. Immunofluorescence was manually quantified using Zen Blue 2.6 software. The percentage of hormone^+^ area per islet was calculated as [(hormone^+^ area/islet area) × 100]. The percentage of beta cells with cytoplasmic proinsulin accumulation was calculated as [(# of insulin^+^ cells with cytoplasmic proinsulin)/(total # of insulin^+^ cells per islet) × 100]. The percentage of beta cells with p21 or gH2AX was calculated as [(# of insulin^+^/p21^+^ cells)/(total # of insulin^+^ cells per islets) × 100] or [(# of insulin^+^/gH2AX^+^ cells)/(total # of insulin^+^ cells per islets) × 100], respectively.

### Statistical Analysis

All statistical analyses were conducted using GraphPad Prism 10.1.2 (GraphPad Software Inc). Specific statistical tests and sample sizes are indicated in the figure legends. For all analyses, p<0.05 was considered statistically significant. Data are presented as mean ± SEM.

## Results

### *In vitro* cisplatin exposure does not affect islet cell viability within 48 hours

We first determined if our *in vitro* cisplatin dosing protocol affected islet cell viability. Isolated male mouse islets were transferred to complete RPMI media containing 10 μM cisplatin or vehicle control. At 6, 24, and 48 hours, intact islets were imaged to visualize morphology, then dispersed into a single cell suspension to measure cell viability via an image-based LIVE/DEAD^TM^ assay (Figure 1A). Islets from both treatment groups appeared to be generally healthy at all timepoints (Figure 1B). Cisplatin exposure did not affect the percentage of PI^+^ (i.e. dead/dying) islet cells at any timepoint (Figure 1C, D). These data indicate that exposure to 10 μM cisplatin for 48 hours was not overtly cytotoxic to mouse islets, so we proceeded with this dosing protocol for all remaining *in vitro* islet experiments.

**Figure 1.**
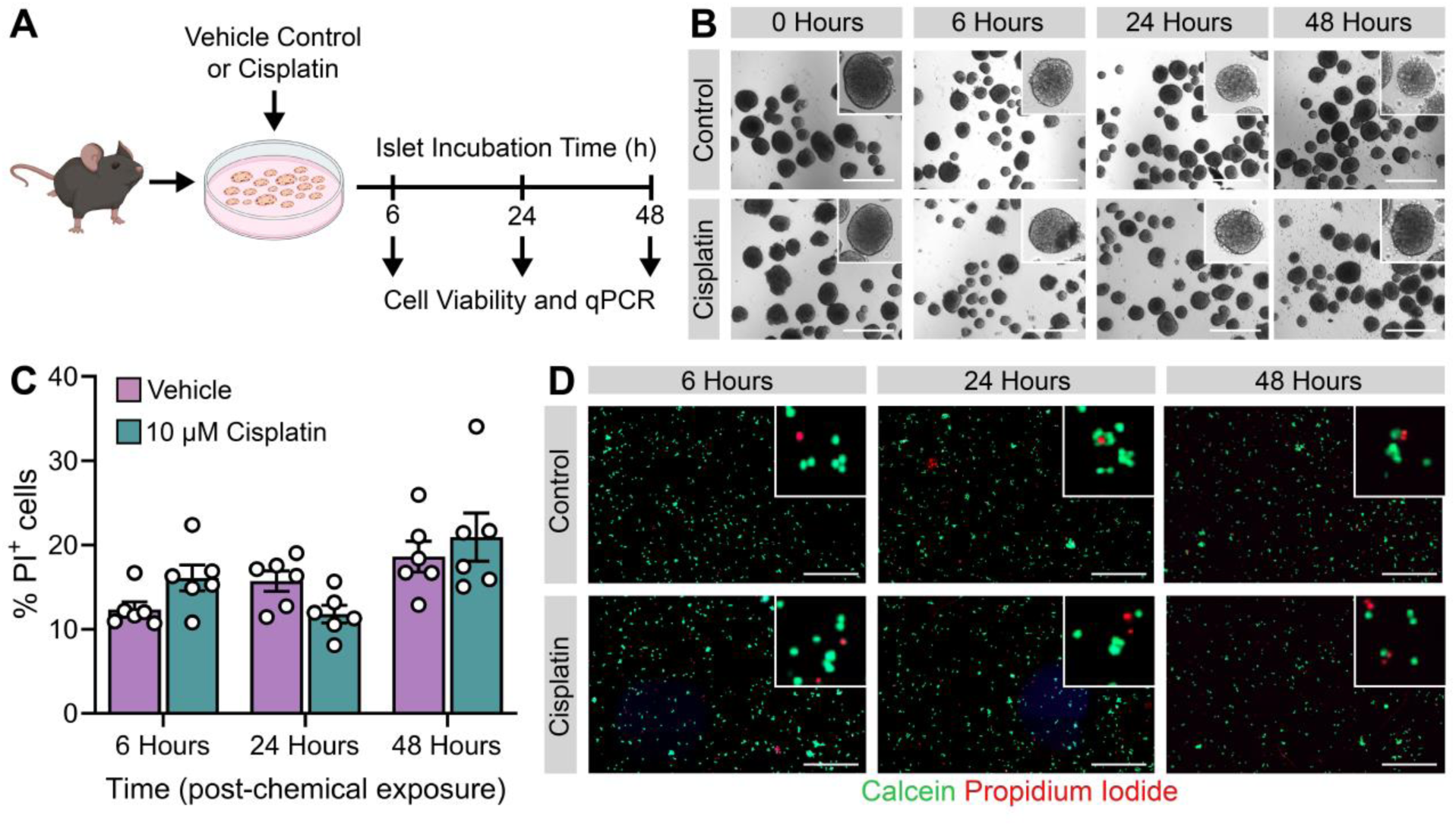
Cisplatin exposure does not cause significant cell death over the course of 48 hours. (A) Schematic summary of *in vitro* exposure protocol. Mouse islets were isolated from male mice and exposed to 10 μM cisplatin or vehicle *in vitro*. A subset of islets was collected after 6, 24, and 48 hours incubation periods and used for live/dead viability assay and qPCR analysis. **(B)** hours after treatment. **(C)** Percentage of cells stained with propidium iodide in a field of view 6, 24, and 48 hours after chemical exposure; n = 6 per treatment group. **(D)** Representative images of live/dead viability assay performed at 6, 24, and 48 hours after chemical exposure. Live cells take up calcein and fluoresce green, dead cells take up propidium iodide and fluoresce red. All scale bars = 500 μm. All data presented as mean ± SEM. Graph was analyzed using a repeated measures two-way ANOVA with Sidak’s multiple comparison test.

### Cisplatin impairs insulin secretion in male mouse islets

To assess the impact of cisplatin on beta cell function, we measured GSIS in vehicle- and cisplatin-exposed mouse islets after 48 hours *in vitro* (Figure 2A). Insulin secretion was first measured using static 1-hour incubations of islets in LG and HG buffer. Cisplatin-exposed islets showed significantly elevated basal insulin secretion under LG conditions (Figure 2B) yet only a small decrease in stimulation index (Figure 2C) and no change in total insulin content (Figure 2D). These data suggest that cisplatin dysregulates insulin secretion without causing pronounced cell loss.

**Figure 2.**
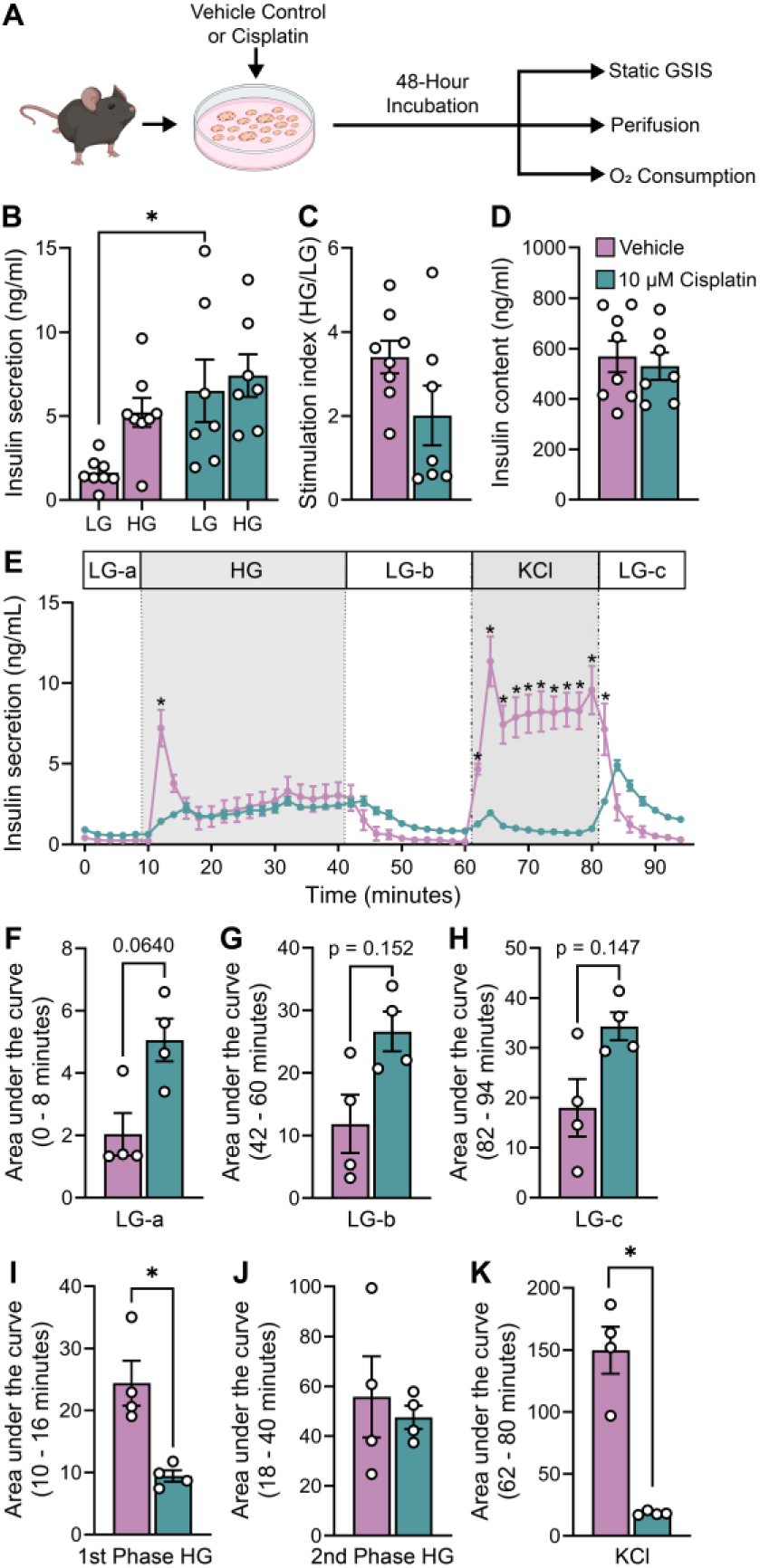
Cisplatin exposure causes dysregulated insulin secretion in primary male mouse islets. **(A)** Schematic summary of *in vitro* exposure protocol. Mouse islets were isolated from male mice and exposed to 10 μM cisplatin or vehicle for 48 hours *in vitro*. **(B)** Static insulin secretion was determined using static GSIS following sequential 1-hour incubations in low glucose (LG; 2.8 mM) and high glucose (HG; 16.7 mM) buffer. **(C)** Islet insulin content was measured following an overnight incubation in acid ethanol. **(D)** The stimulation index was calculated as the ratio of insulin secretion under HG conditions to LG conditions. **(E)** Insulin secretion was measured dynamically every 2 minutes while islets were perfused with LG, HG, and potassium chloride (KCl; 30 mM) buffer. **(F–K)** Area under the curve for islets perfused with **(F–H)** LG buffer, **(I)** HG buffer during first phase insulin secretion, **(J)** HG buffer during second phase insulin secretion, and **(K)** KCl buffer. All data is presented as mean ± SEM. * p<0.05; n= 4-8 per treatment group. The following statistical tests were done: **(B, E)** repeated measures two-way mixed-effects ANOVA with Sidak’s multiple comparison test, **(C–D, F–K)** two-tailed paired t-test.

To assess beta cell function in more detail, we next used perifusion to measure insulin secretion every 2 minutes in cisplatin- and vehicle-exposed islets (Figure 2E). Consistent with the static GSIS results (Figure 2B), cisplatin exposure caused a modest, although not statistically significant, increase in the area under the curve for insulin secretion during all 3 LG incubations (Figure 2E-H). Cisplatin exposure significantly reduced the first phase insulin secretion response to HG (Figure 2E, I) but interestingly did not affect the second phase response to HG (Figure 2E, J). Lastly, KCl-stimulated insulin secretion was nearly abolished in cisplatin-exposed islets (Figure 2E, K) and instead there was a delayed peak in insulin secretion post-KCl (Figure 2E). The perifusion data suggest that cisplatin reduces the sensitivity of islets to glucose, but also impairs the release of insulin, independent of glucose metabolism.

### Cisplatin exposure significantly reduces oxygen consumption in male mouse islets

Mitochondria play a central role in driving glucose-stimulated insulin release, by metabolizing glucose to generate ATP and other metabolic signals. To better understand how cisplatin disrupts insulin secretion (Figure 2B-K), we assessed mitochondrial function in vehicle- and cisplatin-exposed male mouse islets after 48-hours using a Seahorse XFe24 analyzer (Figure 2A). Cisplatin-exposed islets had significantly impaired oxygen consumption compared to vehicle-exposed islets throughout the assay (Figure 3A) and did not robustly increase oxygen consumption when stimulated with HG (Figure 3B). Cisplatin-exposed islets showed a significant or trending reduction in all calculated parameters from the oxygen consumption assay, including basal respiration, maximal respiration, spare respiratory capacity, mitochondrial ATP production, proton leak, and non-mitochondrial respiration compared to vehicle-exposed islets (Figure 3A, C-H). Collectively, these data indicate *in vitro* cisplatin exposure disrupts mitochondrial function in mouse islets, which is consistent with the impaired GSIS in cisplatin-exposed islets (Figure 2).

**Figure 3.**
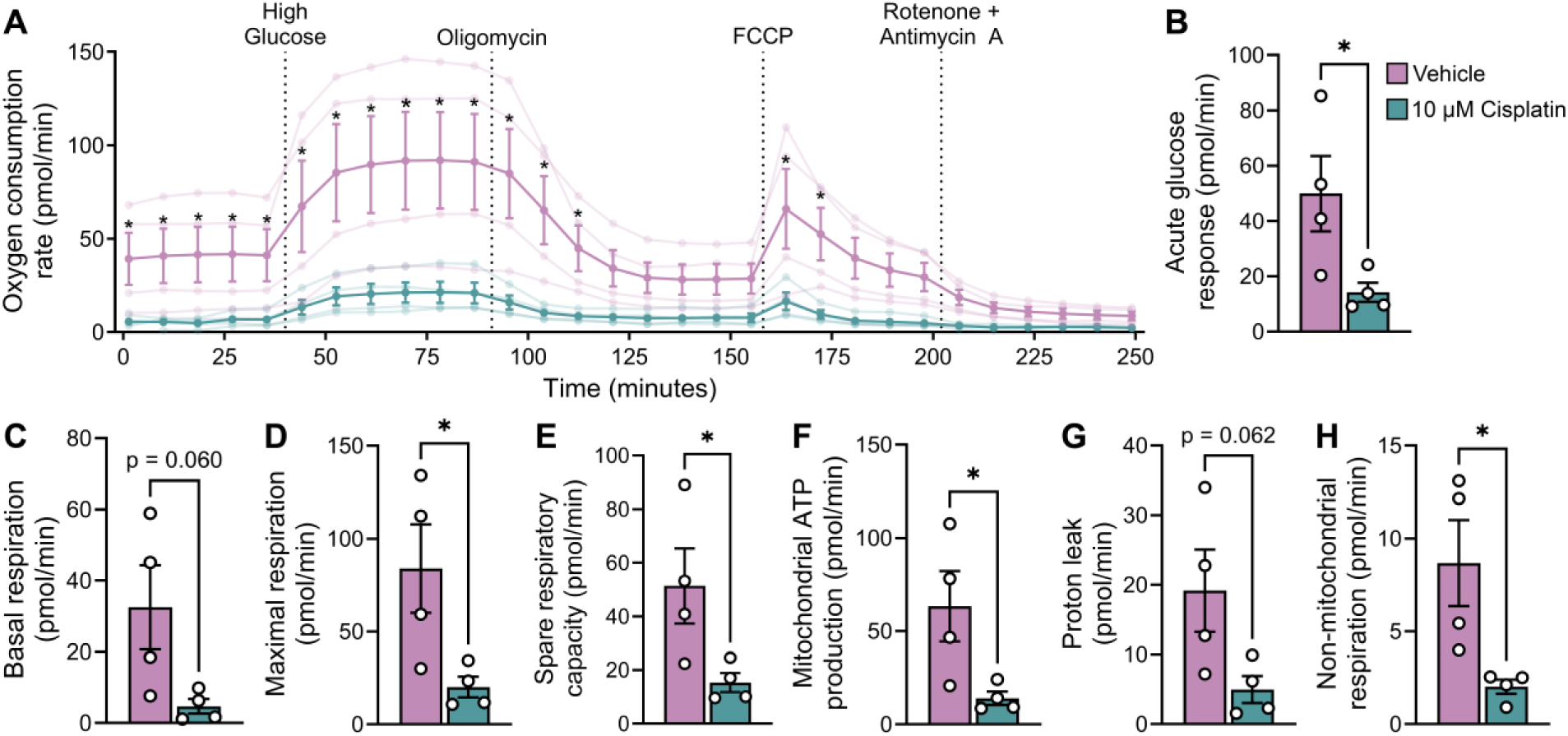
*In vitro* cisplatin exposure impairs mitochondrial function in mouse islets. Mouse islets were isolated from male mice and exposed to 10 μM cisplatin or vehicle for 48 hours *in vitro*. Oxygen consumption rate was measured using a Seahorse XFe24 Analyzer. **(A)** Mouse islets were cultured in low glucose (2.8 mM) media then exposed to serial injections of 16.7 mM glucose, 2.5 μM oligomycin, 3 μM carbonyl cyanide-p-trifluoromethoxyphenylhydrazone (FCCP), and 3 μM combination of rotenone with antimycin A. **(B-H)** Parameters of mitochondrial function including **(B)** acute glucose response, **(C)** basal respiration, **(D)** maximal respiration, **(E)** spare respiratory capacity, **(F)** ATP production from mitochondrial respiration, **(G)** proton leak, and **(H)** non-mitochondrial respiration. All data presented as mean ± SEM. * p<0.05; n=4 per treatment group. The following statistical tests were done: **(A)** repeated measures two-way mixed-effects ANOVA with Sidak’s multiple comparison test, **(B-I)** two-tailed unpaired t-test.

### Cisplatin exposure alters transcription of genes involved in regulating insulin processing, oxidative stress, and apoptosis in mouse islets

We measured expression of key genes related to islet function and cell stress at 6, 24, and 48 hours following vehicle or cisplatin exposure to better understand the temporal effects of cisplatin (Figure 4). Cisplatin did not affect expression of insulin (*Ins1*, *Ins2*) or proprotein convertase genes (*Pcsk1, Pcsk2*) at 6 hours, but reduced expression of these genes at 24 and 48 hours compared to vehicle (Figure 4A-D). The most pronounced effect was on *Pcsk2* expression, which was reduced ∼4-fold at 24 hours and ∼8-fold at 48 hours in cisplatin-exposed islets (Figure 4D). These results suggest cisplatin exposure impairs proinsulin production and processing.

**Figure 4.**
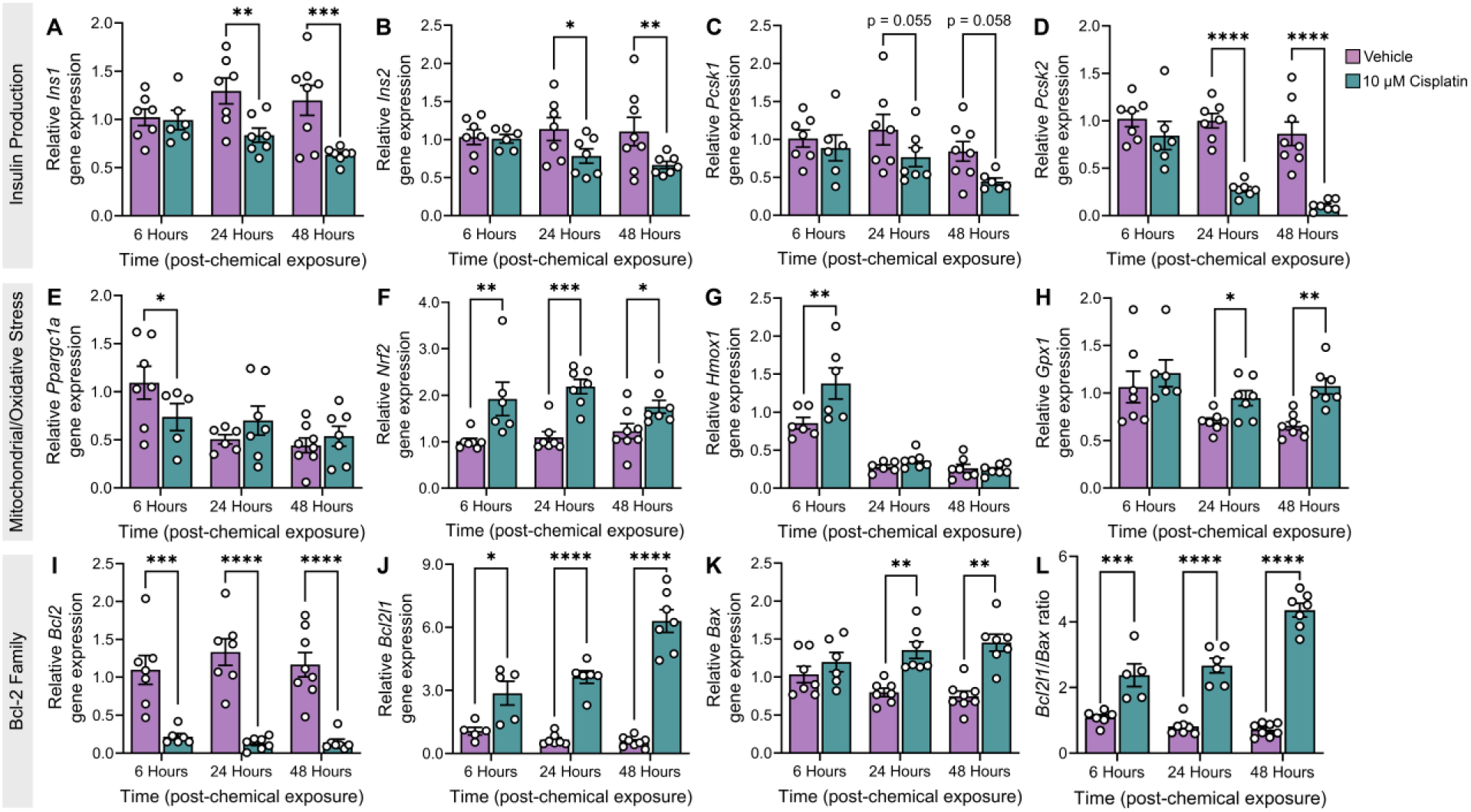
*In vitro* cisplatin exposure alters gene expression of key genes linked to β-cell health and function. Islets were isolated from male mice and exposed to 10 μM cisplatin or vehicle *in vitro*. A subset of islets was collected after 6-, 24-, and 48-hour incubation periods and used qPCR analysis. Expression of **(A)** *Ins1*, **(B)** *Ins2*, **(C)** *Pcsk1*, **(D)** *Pcsk2*, **(E)** *Ppargc1a*, **(F)** *Nrf2*, **(G)** *Hmox1*, **(H)** *Gpx1*, **(I)** *Bcl2*, **(J)** *Bcl2l1*, **(K)** *Bax*, and **(L)** the ratio of *Bcl2l1*-to-*Bax* expression at 3 timepoints relative to 6-hour vehicle control gene expression. All data presented as mean ± SEM. *p<0.05, **p<0.01, ***p<0.001; n=5-7 per treatment group. All graphs were analyzed using a repeated measures two-way mixed-effects ANOVA with Tukey’s multiple comparison test.

*Ppargc1a*—a transcriptional coactivator that regulates mitochondrial function, including respiration and detoxification of ROS(30,31)—was modestly reduced in cisplatin-exposed islets at 6 hours but not 24 or 48 hours (Figure 4E). Cisplatin-exposed islets had ∼2-fold upregulation of *Nrf2*, a marker of oxidative stress, at all 3 timepoints (Figure 4F) and increased expression of *Nrf2* downstream targets *Hmox1* at 6 hours and *Gpx1* at 24 and 48 hours (Figure 4G-H). These changes imply that cisplatin acutely activates oxidative stress responses in mouse islets.

Lastly, we assessed expression of genes in the Bcl-2 family, key players in the intrinsic apoptosis pathway. Cisplatin-exposed islets showed a sustained ∼5-fold downregulation of the anti-apoptotic gene *Bcl2* compared to control islets between 6 to 48 hours post-exposure (Figure 4I), but a progressively increasing upregulation of *Bcl2l1*, another anti-apoptotic gene, throughout the time course (Figure 4J). The pro-apoptotic gene *Bax* was upregulated ∼2-fold in cisplatin-exposed islets at both 24 and 48 hours (Figure 4K). Interestingly, the ratio of *Bcl2l1:Bax* (anti-apoptotic:pro-apoptotic genes) was significantly higher in cisplatin-exposed islets compared to vehicle-exposed islets (Figure 4L). Together these data suggest cisplatin induces the intrinsic apoptosis pathway, but anti-apoptotic gene *Bcl2l1* is being activated to promote a pro-survival phenotype.

### Cisplatin-exposed male mice were hypoglycemic and hypoinsulinemic but did not have altered insulin sensitivity *in vivo*

We next assessed how cisplatin impacts beta cell function *in vivo*. Male mice were exposed to 7 injections of saline (vehicle control) or 2 mg/kg cisplatin every other day for 2 weeks, then tracked for an additional 6 weeks (Figure 5A). Cisplatin exposure caused ∼10% weight loss during the exposure period, leading to a sustained reduction in body weight for the following 4 weeks (Figure 5B). Cisplatin had no effect on fasting blood glucose levels throughout the study (Figure 5C).

**Figure 5.**
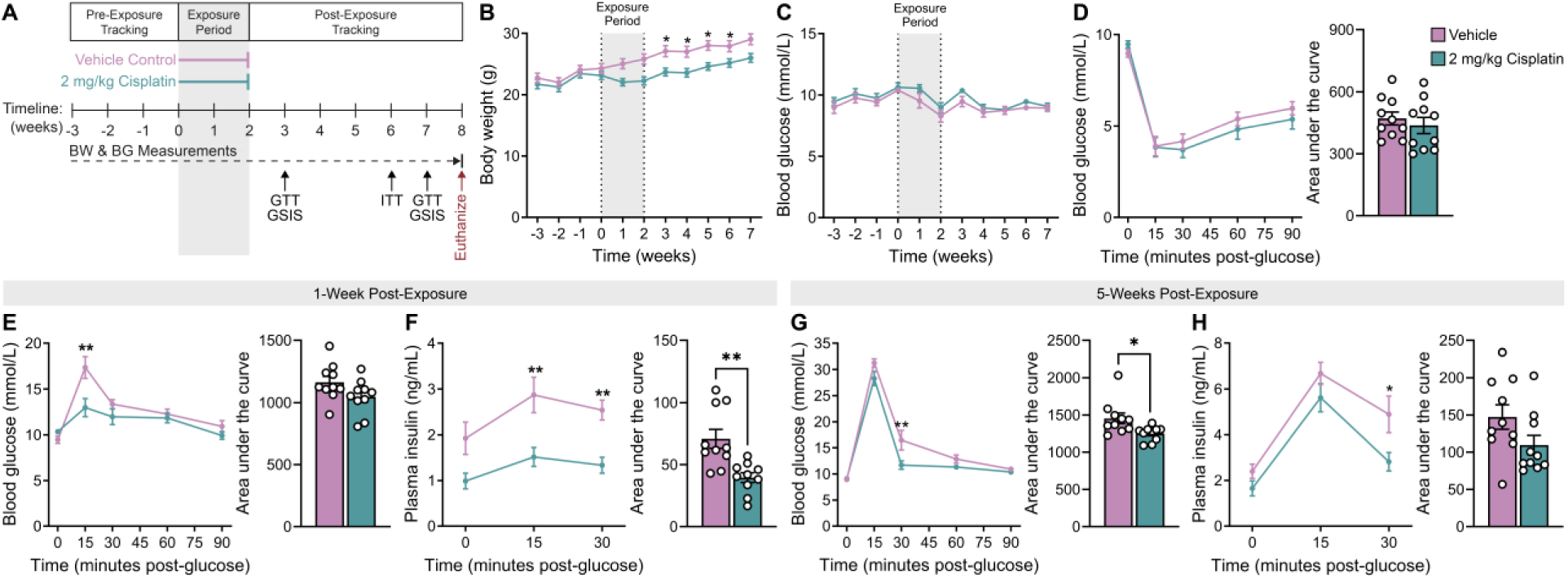
*In vivo* cisplatin exposure impairs glucose tolerance and glucose-stimulated insulin secretion. **(A)** Schematic summary timeline of the study. Male mice were injected with vehicle control or 2 mg/kg cisplatin every other day over the course of 2 weeks, then tracked for 6 weeks following the exposure period. Mice were euthanized 6 weeks post-exposure. BW = body weight, BG = blood glucose, GTT = glucose tolerance test, GSIS = glucose-stimulated insulin secretion, ITT = insulin tolerance test. **(B)** Body weight and **(C)** fasted blood glucose were measured weekly. **(D–H)** Presented as line graphs and area under the curve. **(D)** Blood glucose levels following an ITT 4 weeks post-exposure. **(E, G)** Blood glucose levels and **(F, H)** plasma insulin levels during a GTT **(E, F)** 1-week and **(G, H)** 5 weeks post-exposure. All data presented as mean ± SEM. *p<0.05, **p<0.01; n=10 per treatment group. The following statistical analyses were used: **(B– E, G)** line graphs, repeated measures two-way ANOVA with Sidak’s multiple comparison test; bar graphs, two-tailed unpaired t-test, **(F, H)** line graph, repeated measures two-way mixed-effects ANOVA with Sidak’s multiple comparison test; bar graph, two-tailed unpaired t-test.

Interestingly, during an ipGTT at 1-week post-exposure, cisplatin-exposed mice were modestly hypoglycemic at 15 minutes post-glucose (Figure 5E) but also profoundly hypoinsulinemic at 15- and 30 minutes post-glucose (Figure 5F). An ITT at 4 weeks post-exposure demonstrated that cisplatin-exposed mice had comparable insulin sensitivity to vehicle-exposed controls (Figure 5D). When a second ipGTT was conducted at 5 weeks post-exposure with a higher dose of glucose, cisplatin-exposed mice still had lower blood glucose (Figure 5G) and plasma insulin levels (Figure 5H) compared to vehicle-exposed controls, but only at 30 minutes post-glucose. The diminishing severity of both hypoglycemia and hypoinsulinemia over time (Figure 5E–H) suggest that the effects of cisplatin on metabolic function are lasting but improve with time.

### Cisplatin-exposed mice had increased DNA damage in beta cells and altered proinsulin immunoreactivity

We next assessed islet morphology and other beta cell characteristics in pancreas sections from mice at 6 weeks post-exposure. Cisplatin did not affect average islet size (Figure 6A) or the % of insulin^+^ or glucagon^+^ area per islet (Figure 6B, C). However, there was a significant increase in the proportion of beta cells with cytoplasmic proinsulin accumulation as opposed to the canonical pattern of perinuclear proinsulin immunoreactivity in cisplatin-exposed mice (Figure 6D). There was also a significant increase in the proportion of beta cells expressing the DNA damage markers gH2AX (Figure 6E, H) and p21 (Figure 6F, I).

**Figure 6.**
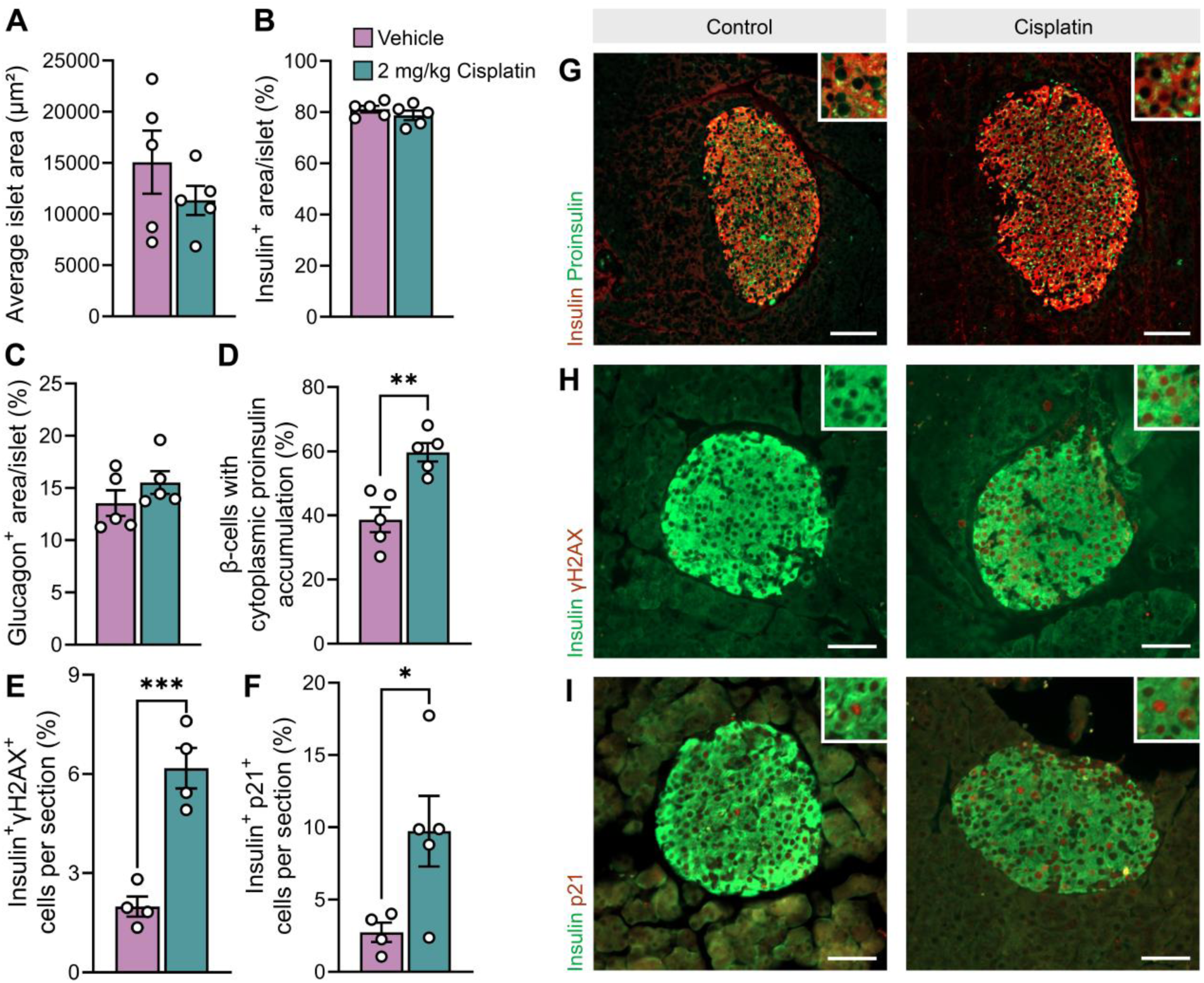
Cisplatin-exposed mice had no change in islet area but increased cytoplasmic proinsulin accumulation and DNA damage markers in β-cells. Whole pancreas was harvested 6 weeks post-exposure period (see Figure 5A for study timeline). **(A)** Average islet size, **(B)** percentage of insulin^+^ area per islet, **(C)** percentage of glucagon^+^ area per islet, **(D)** percentage of β-cells with cytoplasmic proinsulin accumulation, **(E)** percentage of insulin^+^ and gH2AX^+^ cells per mouse, and **(F)** percentage of insulin^+^ and p21^+^ cells per mouse as determined by immunofluorescent staining (n = 5 per group). **(G-I)** Representative images of pancreas sections showing immunofluorescence staining of **(G)** insulin and proinsulin, **(H)** insulin and gH2AX, and **(I)** insulin and p21. Scale bar = 500 μm. All data presented as mean ± SEM. *p<0.05, **p<0.01. All graphs were analyzed using a two-tailed unpaired t-test.

## Discussion

Several studies have reported increased incidence of new onset T2D in cancer survivors following treatment(4,7). Additionally, cancer survivors who received cisplatin treatment had increased prevalence of metabolic syndrome and T2D compared to the general population(15,16). Our research in mice shows that cisplatin exposure profoundly dysregulates insulin secretion and leads to sustained DNA damage in beta cells. Importantly, the adverse effects of cisplatin on insulin secretion were seen both after exposure of mouse islets directly *in vitro* and systemically *in vivo*. These data suggest that cisplatin-induced damage to pancreatic islets may contribute to the long-term risks of metabolic dysregulation in cancer survivors.

Male mouse islets exposed to cisplatin *in vitro* showed higher insulin release under LG conditions, reduced first-phase GSIS, and profoundly suppressed KCl-stimulated insulin release. Basal hyperinsulinemia and the loss of first-phase GSIS are predictive markers for T2D(32,33). Although defective GSIS implies that cisplatin causes metabolic dysfunction, the dampened insulin release in response to KCl stimulation suggests cisplatin alters more than just the metabolic capacity of beta cells. It is possible that the premature hypersecretion of insulin under basal glucose conditions reduces the availability of insulin granules in the readily releasable pool, causing a delayed release of insulin upon both HG and KCl stimulation while the readily releasable pool replenishes itself from the reserve pool(34). Our mouse study confirmed that these beta cell defects translated to islets *in vivo* as well. The significantly reduced plasma insulin levels in cisplatin-exposed mice during a GTT is a key marker of beta cell dysfunction(35). Taken together, our data indicate that both *in vitro* and *in vivo* cisplatin exposure adversely impacts insulin secretion in mouse islets, but the exact mechanism of action remains unclear.

Given that there was no change in total insulin content or % PI^+^ islet cells between treatment groups *in vitro*, and no changes in islet size or hormone^+^ area between groups *in vivo*, we speculate that cisplatin-induced impairments in insulin secretion are not driven by beta cell loss, but rather through intrinsic defects within islet endocrine cells. Canonically, cisplatin forms adducts with mitochondrial DNA and inhibits its replication/transcription(11), so we assessed mitochondrial function in islets as a logical starting point. Indeed, cisplatin-exposed islets had decreased basal oxygen consumption and did not display appropriate changes in their oxygen consumption rate in response to high glucose or electron transport chain modulators. Because the basal oxygen consumption of cisplatin-exposed islets is near maximum respiratory capacity, these islets are unable to accommodate changes in energetic demand as effectively as their vehicle-exposed counterparts. Thus, reduced mitochondrial ATP production likely contributes to the dysregulated first-phase GSIS response in cisplatin-exposed islets, but would not explain the observed defects in KCl-induced insulin secretion. Future research should investigate if cisplatin exposure leads to electrophysiological and exocytotic defects through patch-clamp recordings and Ca^2+^ imaging.

Cisplatin is known to induce mitochondrial ROS production and oxidative stress, both of which can contribute to mitochondrial dysfunction and dysglycemia(13,36). The rapid upregulation of *Nrf2*, a master regulator of oxidative stress response pathways, and its downstream targets confirm that cisplatin is likely increasing oxidative stress in islets. beta cells are particularly sensitive to oxidative stress, as they express relatively low levels of antioxidant enzymes compared to other cells(19). Moreover, the downregulation of *Ppargc1a* in cisplatin-exposed islets may inhibit ROS detoxification(31), further contributing to cisplatin-induced oxidative stress in islets. Further investigation is required to better understand if ROS accumulation is involved in driving cisplatin-induced beta cell dysfunction and to determine if an intervention with antioxidants could protect islets from the adverse effects of cisplatin.

Members of the Bcl-2 family are key regulators of beta cell fate(37).The robust decline in *Bcl2* expression, along with the stark upregulation of *Bax* expression, suggests cisplatin induces the intrinsic apoptosis pathway in islets. However, we did not observe increased PI^+^ cells or reduced insulin content in cisplatin-exposed islets, so we speculate that the upregulation of *Bcl2l1* (and likely other anti-apoptotic factors) contributes to protecting cisplatin-exposed islets from cell death. Fiebig *et al.*(38) found that Bcl-x_L_—the protein product of *Bcl2l1*—is more effective at preventing cell death than Bcl-2 in cells treated with etoposide, another chemotherapeutic agent. Other studies have shown that while upregulation of Bcl-x_L_ prevents cell death, it also impairs mitochondrial function, oxygen consumption, and insulin secretion(39,40); thus, the upregulation of *Bcl2l1* may contribute to the dysregulation in oxygen consumption and insulin release in cisplatin-exposed islets. Interestingly, decreased *Bcl2* expression has also been correlated with elevated levels of basal insulin secretion(41), which supports our findings. These anti-apoptotic Bcl-2 proteins also play a role in beta cell senescence(42,43). We found that cisplatin-exposed mice had increased expression of p21 and gH2AX in beta cells after 5 weeks of recovery following treatment. Taken together, our data suggest that cisplatin exposure may promote a pro-survival phenotype in islets, which could shift cell fate towards senescence rather than apoptosis. Studies using senolytics and small molecule agonists/antagonists of the Bcl-2 family will provide additional insight into the role of senescence in cisplatin-induced beta cell dysfunction.

Defective insulin processing is another important feature of T2D pathogenesis and can contribute to abnormal insulin release patterns(44,45). An elevated proinsulin-to-insulin ratio is often detected in patients with T2D(46). The downregulation of *Ins1, Ins2, Pcsk1,* and *Pcsk2* in cisplatin-exposed mouse islets *in vitro* suggests that cisplatin exposure impairs proinsulin production and processing. Interestingly, cisplatin-exposed mice had increased cytoplasmic proinsulin accumulation, but no significant changes in insulin^+^ area. Taken in context with the observed downregulation of insulin processing genes *in vitro*, we speculate that defects in proinsulin production and processing may contribute to the dysregulated GSIS seen in both cisplatin-exposed islets and mice. Although cisplatin-exposed mice showed signs of recovery from hypoglycemia, the lasting defects in proinsulin processing could have long-term implications.

Cisplatin exposure caused a reduction in body weight along with concurrent hypoglycemia and hypoinsulinemia in mice. Low body weight in humans has been linked to reduced blood glucose levels and lower insulin secretion(47,48). The simultaneous occurrence of hypoinsulinemia and hypoglycemia is not a well understood phenomenon, but has been reported in cases of liver injury in underweight elderly patients(49). A limitation to our study is the temporal gap between the first GTT and ITT; although there was no difference in insulin sensitivity between treatment groups at 4 weeks post-exposure, it is possible there may have been an acute difference in insulin tolerance immediately after the exposure period that was missed. However, cisplatin-exposed mice remained modestly hypoglycemic and hypoinsulinemic the week following the ITT. This indicates insulin sensitivity cannot be the sole cause for the observed metabolic dysfunction. Given the known role of cisplatin in organ toxicity(14), it is likely that injuries to peripheral tissues contribute to impaired glycemic control in cisplatin-exposed mice. Future studies should investigate liver phenotypes, in the context of beta cell function, to better characterize the impact of cisplatin on glycemic control.

Our experiments indicate that cisplatin exposure causes acute defects in beta cell function and may have lasting effects on islet health. Although instances of new-onset diabetes in cancer survivors have been documented(4,8), the cause of this dysglycemia has not been well characterized. By studying the effects of cisplatin exposure on islet function, we are beginning to uncover off-target consequences of cisplatin treatment. Understanding how chemotherapeutic drugs cause beta cell injury is critical to designing targeted interventions that will improve long-term metabolic health outcomes in cancer survivors and reduce their risk of developing T2D post-treatment.

## Abbreviations

BG: Blood glucose
BW: Body weight
FBS: Fetal bovine serum
FCCP: Carbonyl cyanide-p-trifluoromethoxyphenylhydrazone
GSIS: Glucose-stimulated insulin secretion
GTT: Glucose tolerance test
HBSS: Hanks’ balanced salt solution
HG: High glucose
ITT: Insulin tolerance test
Ip: Intraperitoneal
KCl: Potassium chloride
KRBH: Krebs-ringer bicarbonate HEPES buffer
LG: Low glucose
PBS: Phosphate buffer saline
PI: Propidium iodide
qPCR: Quantitative real-time PCR
ROS: Reactive oxygen species
T2D: Type 2 diabetes

## Acknowledgements

We are extremely grateful to Dr. Bruce McKay (Carleton) for his mentorship and guidance for this project. We thank Andrea Smith for her early contributions to this project and Dr. Erin van Zyl for their contributions to manuscript revisions.

## Funding

This research was supported by Diabetes Canada. L.B. is supported by a CIHR CGS-D award and an NSERC-CREATE award on behalf of the CIRTN-R2FIC. J.E.B. is supported by an Ontario Early Researcher Award.

## Duality of interest

The authors declare that there is no duality of interest.

## Contribution statement

L.B., K.S.M., and J.E.B. conceived the experimental design. L.B. and J.E.B. wrote the manuscript. L.B., K.S.M., M.P.H., L.G., K.R.C.R., E.P., E.F., E.E.M., J.A.M, and J.E.B. were involved with acquisition, analysis, and interpretation of data. All authors contributed to manuscript revisions and approved the final version of the article.

## Notes

### Competing Interest Statement

The authors have declared no competing interest.

